# Age-related differences in auditory cortex activity during spoken word recognition

**DOI:** 10.1101/2020.03.05.977306

**Authors:** Chad S. Rogers, Michael S. Jones, Sarah McConkey, Brent Spehar, Kristin J. Van Engen, Mitchell S. Sommers, Jonathan E. Peelle

## Abstract

Understanding spoken words requires the rapid matching of a complex acoustic stimulus with stored lexical representations. The degree to which the brain networks supporting spoken word recognition are affected by adult aging remains poorly understood. In the current study we used fMRI to measure the brain responses to spoken words in two conditions: an attentive listening condition, in which no response was required, and a repetition task. Listeners were 29 young adults (aged 19–30 years) and 32 older adults (aged 65–81 years) without self-reported hearing difficulty. We found largely similar patterns of activity during word perception for both young and older adults, centered on bilateral superior temporal gyrus. As expected, the repetition condition resulted in significantly more activity in areas related to motor planning and execution (including premotor cortex and supplemental motor area) compared to the attentive listening condition. Importantly, however, older adults showed significantly less activity in probabilistically-defined auditory cortex than young adults when listening to individual words in both the attentive listening and repetition tasks. Age differences in auditory cortex activity were seen selectively for words (no age differences were present for 1-channel vocoded speech, used as a control condition), and could not be easily explained by accuracy on the task, movement in the scanner, or hearing sensitivity (available on a subset of participants). These findings indicate largely similar patterns of brain activity for young and older adults when listening to words in quiet, but suggest less recruitment of auditory cortex by the older adults.

## Introduction

Understanding spoken words requires mapping complex acoustic signals to a listener’s stored lexical representations. Evidence from neuropsychology and cognitive neuroscience provides increasingly converging evidence about the roles of bilateral temporal cortex (particularly superior temporal gyrus and middle temporal gyrus) in processing speech acoustics and recognizing single words (Binder et al., 2000; Hickok and Poeppel, 2007; Peelle et al., 2010a). However, the degree to which the networks supporting spoken word recognition in older adults is still unclear. The goals of the current study were to test whether young and older adults relied on different brain networks during successful spoken word recognition, and whether any age differences were related to the specific task.

Important themes when considering older adults’ language processing include the degree to which linguistic processing is preserved, and whether older adults may adopt different strategies when understanding language compared to young adults (Wingfield and Stine-Morrow, 2000; Peelle, 2019). Particularly important for spoken word recognition is that adult aging frequently brings changes to both hearing sensitivity (Peelle and Wingfield, 2016) and cognitive ability (Park et al., 2002). Thus, it is not surprising that older adults’ spoken word perception differs from that of young adults, particularly in the presence of background noise (Humes, 1996). Older adults tend to take longer to recognize words (Wingfield et al., 1991; Lash et al., 2013), make more recognition errors than young adults, and show increased sensitivity to factors such as the number of phonological neighbors (competitors) associated with a given target word (Sommers and Danielson, 1999). An open question centers on the brain networks on which older adults rely during spoken word recognition. Of particular interest is whether additional regions may be recruited to support successful recognition compared to those engaged by young adults.

A number of studies have investigated neural activity during older adults’ speech processing in noise or other acoustic degradation, using an assortment of tasks and testing participants with different levels of hearing (Hwang et al., 2007; Wong et al., 2009; Bilodeau-Mercure et al., 2015; Manan et al., 2015, 2017). Harris and colleagues (2009), for example, examined spoken word recognition in young and older adults. They varied the intelligibility of the target items using low-pass filtering of the acoustic signal. During scanning, participants were asked to repeat back the word they heard. The authors found increased activity in regions associated with word processing, including auditory cortex and premotor cortex, when words were more intelligible; these intelligibility-related changes did not statistically differ between young and older adults. Older adults did show more activation in the anterior cingulate cortex and supplemental motor area than the young adults did, suggesting a possible increase in top-down executive control.

Age differences in speech understanding have also been studied in the context of sentence comprehension. One common finding is that during successful sentence processing, older adults show additional activity compared to young adults (for example, in contralateral homologs to regions seen in young adults, or in regions beyond the network activated by young adults) (Peelle et al., 2010b; Tyler et al., 2010). These findings have been interpreted in a compensation framework in which older adults are less efficient using a core speech network and need to recruit additional regions to support successful comprehension (Wingfield and Grossman, 2006). However, at least some of this additional activity has been shown to be related to the tasks performed by participants in the scanner, which frequently contain metalinguistic decisions not required during everyday conversation (Davis et al., 2014). Thus, it may be that core language computations are well-preserved in aging (Shafto and Tyler, 2014; Campbell et al., 2016).

The role of executive attention in older adults’ spoken word recognition has also been of significant interest. Listening to speech that is acoustically degraded can result in perception errors, after which listeners must re-engage attention systems to support successful listening. The cingulo-opercular network, an executive attention network (Power and Petersen, 2013; Neta et al., 2015), shows increased activity following perception errors (similar to error-related activity in other domains). Crucially, when listening to spoken words in background noise, increased cingulo-opercular activity following one trial is associated with recognition success on the following trial (Vaden et al., 2013, 2016), consistent with a role in maintaining task-related attention (Eckert et al., 2009).

An important challenge when considering the performance of listeners with hearing loss is that words may not be equally intelligible to all listeners. A common measure of accuracy in spoken word recognition is to ask listeners to repeat each word after hearing it; however, this type of task requires motor responses, which may obscure activations related to speech perception and increase participant motion in the scanner (Gracco et al., 2005). In addition, differences in the brain regions coordinating speech production in older adults (Bilodeau-Mercure and Tremblay, 2016; Tremblay et al., 2017) may interfere with clear measurements of activity during perception and recognition. The degree to which motor effects resulting from word repetition may obscure activity related to speech perception is unclear. In sentence processing tasks, task effects can be significant (Davis et al., 2014), and if not accounted for may obscure what are actually consistent patterns of language-related activity across the lifespan (Campbell et al., 2016).

In the current study we investigated spoken word processing in young and older adult listeners in the absence of background noise. We compared paradigms requiring words to be repeated with “attentive listening” (no motor response required). Our interest is, first, whether age differences exist in the brain networks supporting spoken word recognition, and second, whether these differences are affected by the choice of task. Thus, our primary analyses will focus on activity seen for words (greater than noise) in the experimental conditions.

The influence of psycholinguistic factors on spoken word recognition has long been appreciated. In a secondary set of analyses, we will investigate whether word frequency or phonological neighborhood density modulate activity during spoken word recognition. Although behavioral and electrophysiological studies suggest that high frequency words are processed more quickly than low frequency words, the degree to which this might be captured in fMRI is unclear. Similarly, although neighborhood density effects are widely reported in behavioral studies (with words from dense neighborhoods typically being more difficult to process), the degree to which lexical competition effects may differ with age is unclear.

## Method

Stimuli, data, and analysis scripts are available from https://osf.io/vmzag/.

### Participants

We recruited two groups of participants (young and older adults) for this study. The young adults were 29 self-reported healthy, right-handed adults, aged 19–30 years (M = 23.8, SD = 2.9, 19 female), and were recruited via the Washington University in St. Louis Department of Psychological and Brain Sciences Subject Pool. Older adult participants were 32 self-reported healthy, right-handed adults, aged 65–81 years (M = 71.0, SD = 5.0, 17 female). All participants self-reported themselves to be native speakers of American English with no history of neurological difficulty, and with normal hearing (and no history of diagnosed hearing problem). Participants were compensated for their participation, and all provided informed consent commensurate with practices approved by the Washington University in St. Louis Institutional Review Board.

Audiograms were collected on a subset of 8 young and 9 older participants using pure-tone audiometry (**Figure 1a**). We summarized hearing ability using a better-ear pure tone average (PTA) at 1, 2, and 4 kHz. PTAs in participants’ better hearing ears ranged from −3.33 to 8.33 dB HL in young adults (M = 2.92, SD = 4.15), and 8.33 to 23.3 dB HL in older adults (M = 23.3, SD = 9.17).

**Figure 1.**
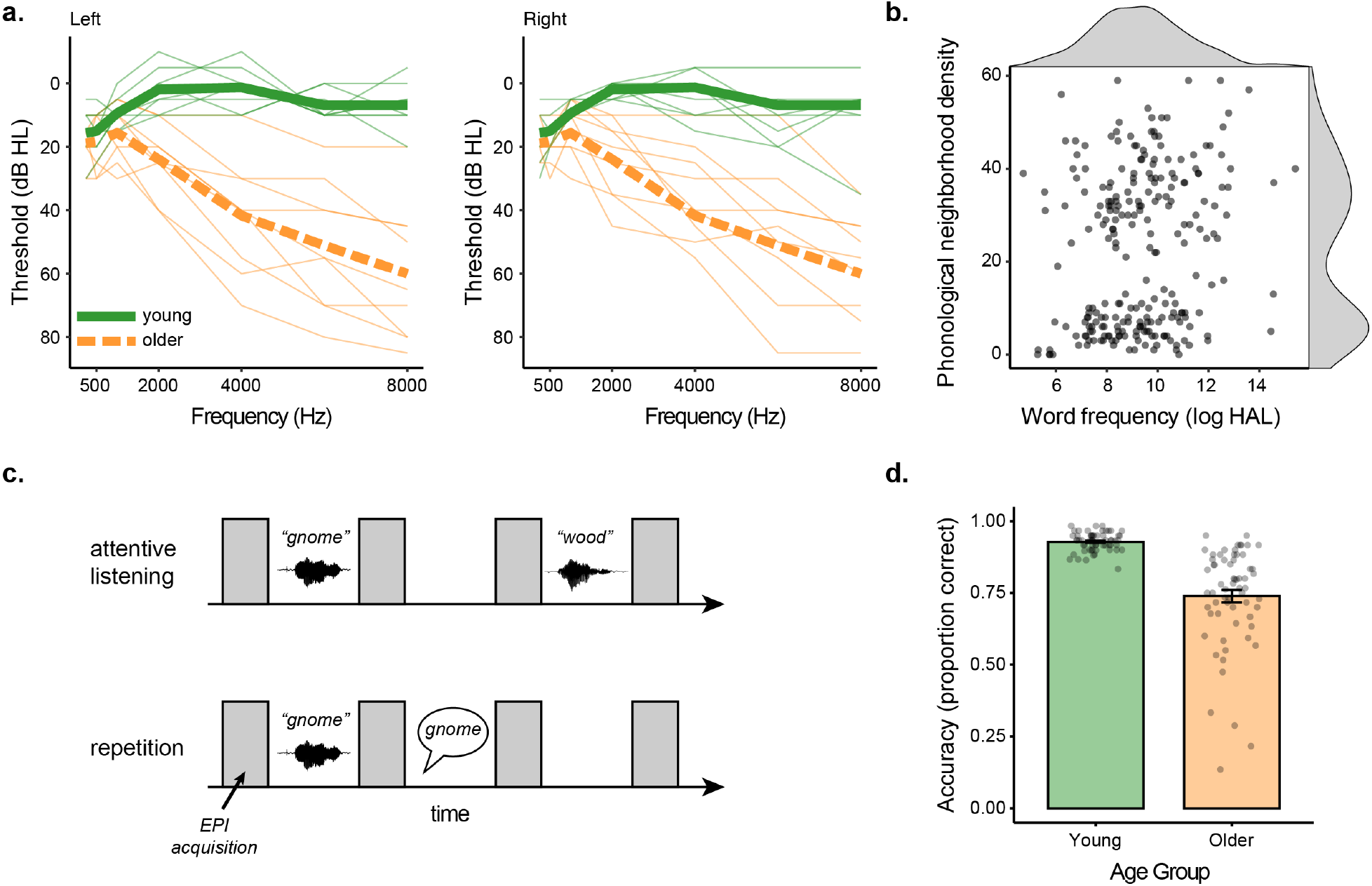
Experiment overview. **a.** Audiograms for the subset of participants on whom hearing was available for left and right ears. Individual participants are shown in thin lines, group means in thick lines. **b.** Frequency of occurrence and phonological neighborhood density for the 240 experimental items. **c.** Task design for attentive listening and word repetition tasks. **d.** Behavioral accuracy for the repetition condition for young and older adults.

### Materials

Stimuli for this study were 375 monosyllabic consonant-vowel-consonant (CVC) words. The auditory stimuli were recorded at 48 kHz using a 16-bit Digital-to-Analog converter with an Audio Technica 2045 microphone in a quiet room. Words were spoken by a female speaker with a standard American dialect. Root-mean-square amplitude of the stimuli was equated.

Out of the full set of words, 75 words were vocoded using a single channel with white noise as a carrier signal (Shannon et al., 1995) using *jp_vocode.m* from http://github.com/jpeelle/jp_matlab. These stimuli were used for an unintelligible baseline “noise” condition. The remaining 300 words were divided into five lists of 60 words, using MATCH software (Van Casteren and Davis, 2007), and were balanced for word frequency (as measured by the log of the Hyperspace Analogue to Language (HAL) dataset), orthographic length, concreteness (Brysbaert et al., 2014), and familiarity (Balota et al., 2007). Distribution of word frequency and phonological neighborhood density are shown in **Figure 1b**.

One of these lists was combined with 15 of the noise vocoded words and used for word repetition task practice outside of the scanner. The remaining four lists of 60 words served as the critical items inside the scanner, with half of the lists used for passive listening (120 total words) and the other half for word repetition (120 total words). Word lists were counterbalanced such that each word was presented in both “listen” and “repeat” conditions across participants.

### Procedure

Prior to scanning, participants were taken to a quiet room.^1^ During that time participants provided informed consent, completed demographic questionnaires, and a subset had their hearing tested using a calibrated Maico MA40 portable audiometer (Maico Diagnostics, Inc., Eden Prairie MN) by an audiologist-trained researcher.

Participants were then instructed for the two tasks they would perform in the scanner: attentive listening and word repetition. During attentive listening, participants were asked to stay alert, still, and keep their eyes focused on a fixation cross while listening to a sequence of auditory sounds including words, silence, and noise (single-channel noise vocoded words). During word repetition, participants were asked to do the same as in passive listening, with the addition of repeating the word they just heard aloud. Participants were instructed to repeat the words following the volume acquisition after each word (**Figure 1c**). Participants were told to give their best guess if they could not understand a word. Participants practiced a simulation of the word repetition task until the experimenter was confident that the participant understood the pacing and the nature of the task. Sound levels were adjusted to achieve audible presentations at the beginning of the study and thereafter not adjusted.

Functional MRI scanning took place over the course of four scanning blocks, where participants alternated between blocks of passive listening and word repetition (**Figure 1c**). The order of blocks was counterbalanced such that participants were equally likely to begin with a word repetition or attentive listening block. During word repetition, participants’ spoken responses were recorded using an in-bore Fibersound optical microphone. These responses were scored for accuracy offline by a research assistant (**Figure 1d**).

### MRI data acquisition and processing

MRI data are available from https://openneuro.org/datasets/ds002382 (Poldrack et al., 2013). MRI data were acquired using a Siemens Prisma scanner (Siemens Medical Systems) at 3 T equipped with a 32-channel head coil. Scan sequences began with a T1-weighted structural volume using an MPRAGE sequence [repetition time (TR) = 2.4 s, echo time (TE) = 2.2 ms, flip angle = 8°, 300 × 320 matrix, voxel size = 0.8 mm isotropic]. Blood oxygenation level-dependent (BOLD) functional MRI images were acquired using a multiband echo planar imaging sequence (Feinberg et al., 2010) [TR = 3.07 s, TA = 0.770 s, TE = 37 ms, flip angle = 37°^2^, voxel size = 2 mm isotropic, multiband factor = 8]. We used a sparse imaging design in which there was an 2.3 second delay between scanning acquisitions and the TR was longer than the acquisition time to allow for minimal scanning noise during stimulus presentation and audio recording of participant responses (Edmister et al., 1999; Hall et al., 1999).

Analysis of the MRI data was performed using Automatic Analysis version 5.4.0 (Cusack et al., 2015) (RRID:SCR_003560) which scripted a combination of SPM12 (Wellcome Trust Centre for Neuroimaging) version 7487 (RRID:SCR_007037) and FSL (FMRIB Analysis Group; Jenkinson et al., 2012) version 6.0.1 (RRID:SCR_002823).

Data were realigned using rigid-body image registration, and functional data were co-registered with the bias-corrected T1-weighted structural image. Spatial and functional images were normalized to MNI space using a unified segmentation approach (Ashburner and Friston, 2005), and resampled to 2 mm. Finally, the functional data were smoothed using an 8 mm FWHM Gaussian kernel.

For the attentive listening condition, we did not have measures of accuracy, so we analyzed all trials. For the repetition condition, we analyzed only trials associated with correct responses. For both tasks, we modeled the noise condition in addition to words. Finally, we included three parametric modulators for word events: word frequency, phonological neighborhood density, and their interaction. To avoid order effects (Mumford et al., 2015), these were not orthogonalized.

Motion effects were of particular importance given that participants were speaking during the repetition condition. To mitigate the effects of motion, we used a thresholding approach in which high motion frames were individually modeled for each subject using a delta function in the GLM (see e.g. Siegel et al., 2014). Motion was quantified using framewise displacement (FD), calculated from the 6 motion parameters estimated during realignment assuming the head is a sphere having a radius of 50 mm (Power et al., 2012). We then chose an FD threshold (0.561) that we used for all participants. Our rationale was that some participants move more, and thus produce worse data; we therefore wanted to use a single threshold for all participants, resulting in more data exclusion from high-motion participants. This threshold resulted in 2.2– 19.4% (M = 6.21, SD = 4.45) data exclusion for the young adults and 2.8–58.4% (M = 22.6, SD = 15.3) data exclusion for the older adults. For each frame exceeding this threshold, we added a column to that participant’s design matrix consisting of a delta function at the time point in question, which effectively excludes the variance of that frame from the model.

Contrast images from single subject analyses were analyzed at the second level using permutation testing (FSL *randomise;* 5000 permutations) with a cluster-forming threshold of p < .001 (uncorrected) and results corrected for multiple comparisons based on cluster extent (p < .05). Images (contrast images and unthresholded t maps) are available from https://identifiers.org/neurovault.collection:6735 (Gorgolewski et al., 2015). Anatomical localization was performed using converging evidence from author experience (Devlin and Poldrack, 2007) viewing statistical maps overlaid in MRIcrogl (Rorden and Brett, 2000), supplemented by atlas labels (Tzourio-Mazoyer et al., 2002).

For region of interest (ROI) analysis of primary auditory cortex, we used probabilistic maps based on postmortem human histological staining (Morosan et al., 2001) available in the SPM Anatomy toolbox (Eickhoff et al., 2005) (RRID:SCR_013273). We created a binary mask for regions Te1.0 and Te1.1 then extracted parameter estimates for noise and word contrasts for the attentive listening and repetition conditions from each participant’s first-level analyses by averaging over all voxels in each ROI (left auditory, right auditory).

Outputs from analysis stages used for quality control are available from https://osf.io/vmzag/in the `aa_report` folder.

## Results

### Behavioral data

We analyzed the accuracy data using a linear mixed effects analysis, implemented using the lme4 and lmerTest packages in R version 3.6.2 (RRID:SCR_001905). Because trial-level accuracy data was binary we used logistic regression. We first tested for age differences using a model that included age group as a fixed factor and subject as a random factor:

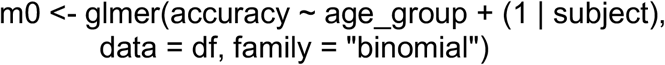

The estimate of age_group was −1.4929 (SE = 0.1902), p = 4.24e-15, consistent with a main effect of age (older adults performing more poorly). Because of ceiling effects and lack of variability in the young adult data we ran an additional analysis only in the older adults, using a model that included neighborhood density and word frequency as fixed factors, and item and subject as random factors:

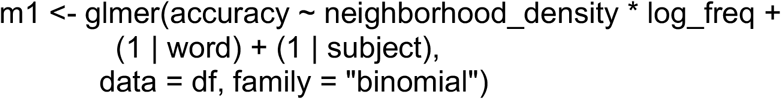

The model failed to converge when including a more complex random effect structure. The results of this model are shown in Table 1. There were no significant effects of neighborhood density or word frequency in the accuracy data.

**Table 1.**
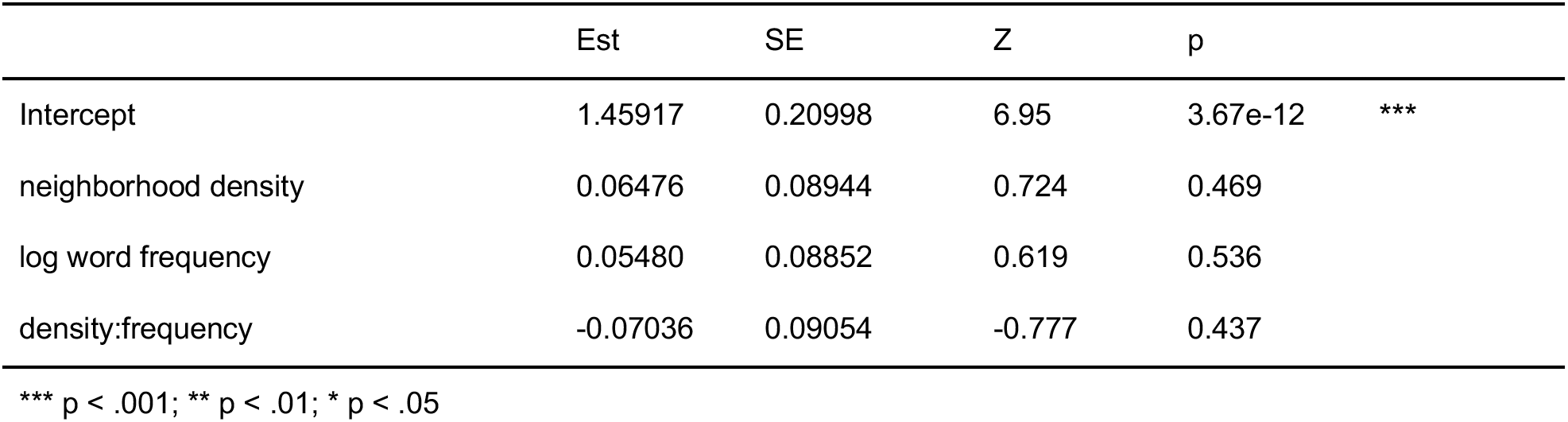
Fixed effects results for accuracy data model

|  | Est | SE | Z | p |  |
| --- | --- | --- | --- | --- | --- |
| Intercept | 1.45917 | 0.20998 | 6.95 | 3.67e-12 | *** |
| neighborhood density | 0.06476 | 0.08944 | 0.724 | 0.469 |  |
| log word frequency | 0.05480 | 0.08852 | 0.619 | 0.536 |  |
| density:frequency | -0.07036 | 0.09054 | -0.777 | 0.437 |  |

\*\*\* $p < .001$ ; \*\* $p < .01$ ; \* $p < .05$

There are many reasons a participant might make an incorrect response, and our primary interest is on the processes supporting successful comprehension. Thus, for the fMRI analyses, we restricted our analyses to correct trials only.

### Imaging Data

We began by looking at activity in auditory cortex, followed by a whole-brain analyses. Activity in left and right auditory cortex for noise and word conditions for young and older adults is shown in **Figure 2a**. We analyzed these data using a linear mixed effects analysis, implemented using the *lme4* and *lmerTest* packages in R version 3.6.2 (RRID:SCR_001905). The full model included task (listen, repeat), stimulus (word, noise), hemisphere (left, right), age group (young, older), and accuracy on the repetition task as fixed factors, with subject, stimulus type, and task as random factors:

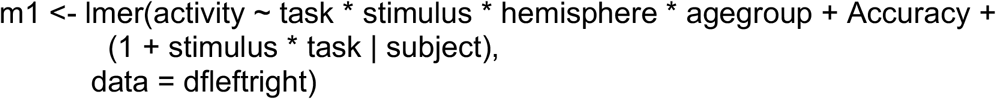

**Figure 2.**
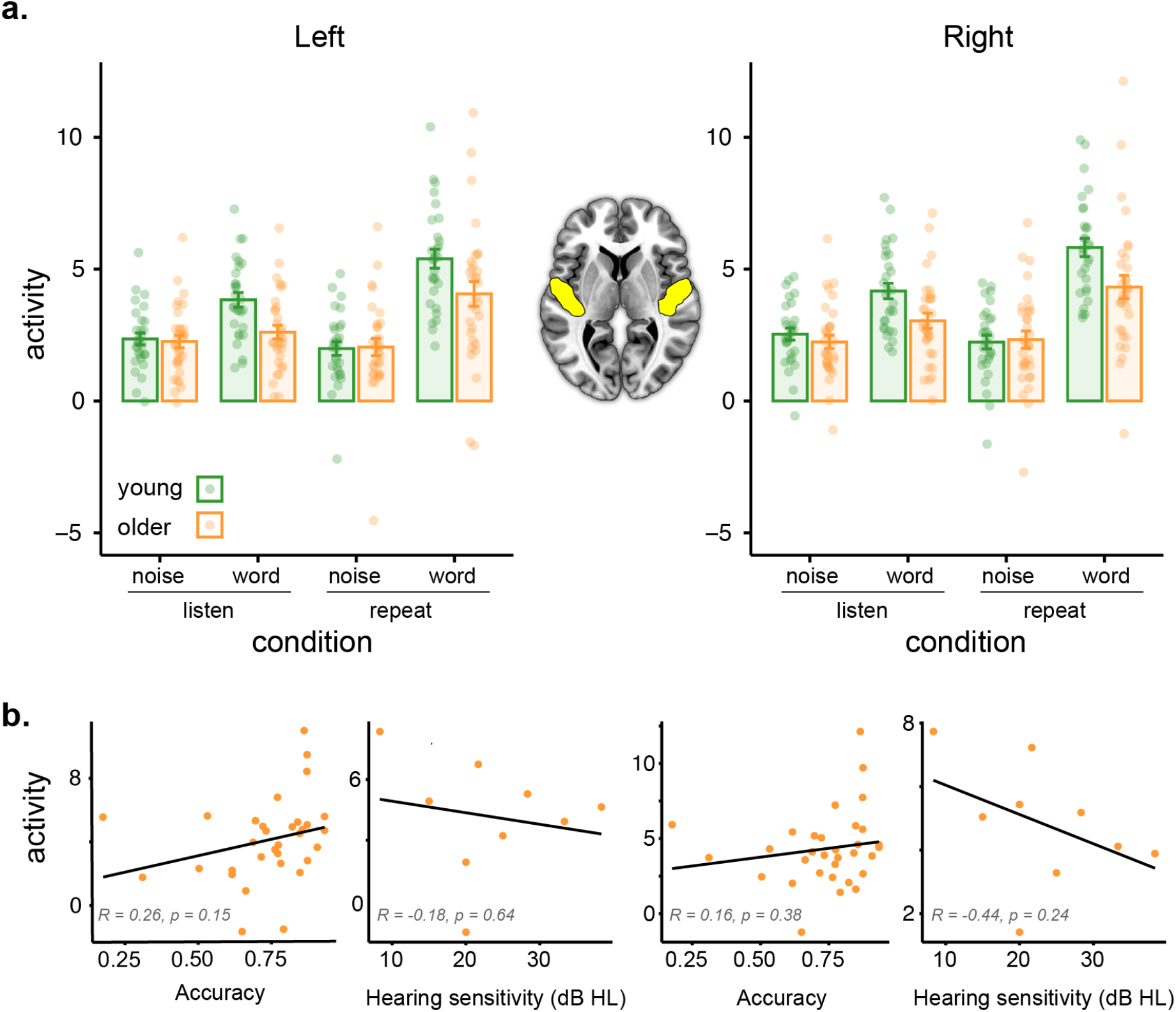
Activity in auditory cortex regions of interest. **a.** Activity (parameter estimates, arbitrary units) for left and right auditory cortex as a function of age group and task. Participants are indicated by individual dots; mean ± standard error indicated by error bars. **b.** Activity for left and right auditory cortex during the word repetition task in older adults as a function of accuracy and hearing (hearing only available in a subset of participants).

When including hemisphere as an additional random factor the model failed to converge, and as our main interests lay elsewhere we settled on the above model.

Full model results are shown in Table 2. P-values were obtained from the *lmerTest* package using Satterthwaites’s method for degrees-of-freedom and t-statistics. We found significant interactions between task and stimulus, consistent with a greater degree of activation for words relative to noise in the repetition task compared to the attentive listening task. Importantly, there was a significant interaction between stimulus and age group, consistent with greater age differences for words relative to noise. We verified this with follow-up t-tests, collapsing over hemisphere, which showed a significant difference in activity between young and older adults for words [attentive listening: t(57.973) = 3.1428, p = 0.002636; repetition: t(56.619) = 2.5583, p = 0.01322] but not for noise [attentive listening: t(58.559) = 0.66361, p = 0.5095; repetition: t(57.241) = 0.19028, p = 0.8498]. None of the other main effects or interactions were significant.

**Table 2.** Fixed effects results for auditory cortex model.

|  | Est | SE | df | t | p |  |
| --- | --- | --- | --- | --- | --- | --- |
| Intercept | 1.76976 | 0.84161 | 60.91 | 2.103 | 0.03962 | * |
| task | -0.21168 | 0.32741 | 85.14 | -0.647 | 0.51966 |  |
| stimulus | 0.35568 | 0.32741 | 125.70 | 1.518 | 0.13156 |  |
| hemisphere | -0.01502 | 0.19203 | 236.00 | -0.078 | 0.93771 |  |
| age | -0.02927 | 0.39170 | 82.77 | -0.075 | 0.94061 |  |
| accuracy | 0.65940 | 1.09603 | 58.00 | 0.602 | 0.54977 |  |
| task:stimulus | 1.66089 | 0.37515 | 105.02 | 4.427 | 2.34e-05 | *** |
| task:hemisphere | 0.29972 | 0.27157 | 236.00 | 1.104 | 0.27087 |  |
| stimulus:hemisphere | 0.44485 | 0.27157 | 236.00 | 1.638 | 0.10274 |  |
| task:age | -0.15187 | 0.47484 | 85.14 | -0.320 | 0.74988 |  |
| stimulus:age | 1.13313 | 0.33986 | 125.70 | 3.334 | 0.00112 | ** |
| hemisphere:age | 0.19873 | 0.27851 | 236.00 | 0.714 | 0.47620 |  |
| task:stimulus:hemisphere | -0.46865 | 0.38406 | 236.00 | -1.22 | 0.22359 |  |
| task:stimulus:age | 0.25428 | 0.54409 | 105.02 | 0.467 | 0.64122 |  |
| task:hemisphere:age | -0.23793 | 0.39387 | 236.00 | -0.604 | 0.54636 |  |
| stimulus:hemisphere:age | -0.29720 | 0.39387 | 236.00 | -0.755 | 0.45126 |  |
| task:stimulus:hemisphere:age | 0.50413 | 0.55701 | 236.00 | 0.905 | 0.36635 |  |

\*\*\* $p < .001$ ; \*\* $p < .01$ ; \* $p < .05$

To explore the possible contribution of other factors to older adults’ reduced activity in auditory cortex, we conducted a series of exploratory correlation analyses with accuracy, hearing, and movement parameters from fMRI (median FD). None of these analyses showed significant correlations with auditory cortex activity. Correlations for accuracy and hearing in the word repetition task are shown in **Figure 2b**. Overall, we interpret these results as being consistent with less auditory activity for the older adults during spoken word perception (but not during our nonspeech control condition).

To complement the ROI analyses, we next performed whole-brain analyses for all conditions of interest. Activity for word perception for the attentive listening condition (greater than the noise baseline) is shown in **Figure 3**(with maxima listed in **Tables 3– 5**). As expected, both young and older adults showed significant activity in bilateral superior temporal cortex. Young adults showed significantly stronger activity in superior temporal cortex near auditory cortex. There were no regions in which older adults showed greater activity than young adults.

**Figure 3.**
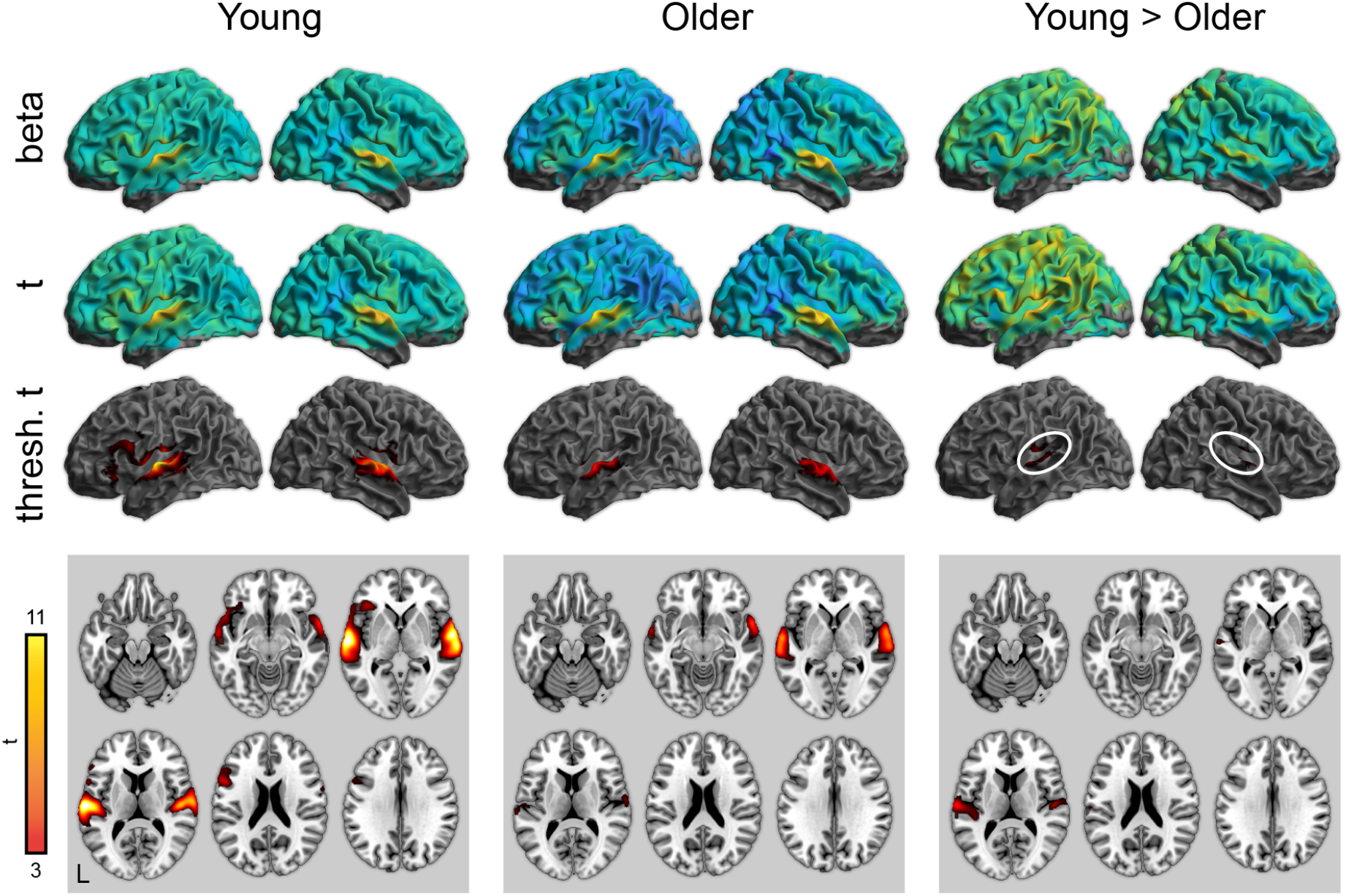
Whole-brain activity for the attentive listening condition. **Top:** Unthresholded parameter estimates. **Middle:** Unthresholded t maps. **Bottom:** Thresholded t maps (p < .05, cluster corrected). White ovals highlight left and right auditory cortex.

**Table 3.** Peak activations for attentive listening condition greater than noise, young adults

| Region | Size (μl) | t score | Coordinates | | |
| --- | --- | --- | --- | --- | --- |
|  |  |  | x | y | z |
| Left superior temporal gyrus | 38,936 | 13.6 | -62 | -16 | 6 |
| Left inferior frontal gyrus |  | 5.44 | -52 | 12 | 20 |
| Left inferior frontal gyrus |  | 5.08 | -38 | 28 | 0 |
| Left inferior frontal gyrus |  | 5.0 | -50 | 20 | -6 |
| Left inferior frontal gyrus |  | 4.75 | -52 | 28 | 16 |
| Left insula |  | 4.08 | -30 | 24 | -4 |
| Left inferior frontal gyrus |  | 3.89 | -34 | 32 | -14 |
| Left inferior frontal gyrus |  | 3.7 | -38 | 26 | -14 |
| Left precentral gyrus |  | 3.46 | -60 | 2 | 26 |
| Right superior temporal gyrus | 23,432 | 12.4 | 60 | -8 | 4 |
| Right superior temporal gyrus |  | 10.2 | 66 | -18 | 0 |
| Right superior temporal gyrus |  | 4.07 | 48 | 12 | -12 |
| Supplemental motor area | 9,144 | 5.39 | -2 | 6 | 56 |
| Supplemental motor area |  | 4.33 | 0 | -4 | 64 |
| Supplemental motor area |  | 4.01 | -8 | 0 | 76 |
| Left paracentral lobule |  | 3.93 | -4 | -28 | 78 |
| Left paracentral lobule |  | 3.84 | -4 | -16 | 80 |
| Left postcentral gyrus |  | 3.61 | -18 | -28 | 76 |
| Supplemental motor area |  | 3.6 | 0 | 8 | 72 |

**Table 4.** Peak activations for attentive listening condition greater than noise, older adults

| Region | Size (μl) | t score | Coordinates | | |
| --- | --- | --- | --- | --- | --- |
|  |  |  | x | y | z |
| Right superior temporal gyrus | 11,152 | 7.99 | 62 | -8 | 2 |
| Right superior temporal gyrus |  | 7.78 | 60 | 0 | -4 |
| Right superior temporal gyrus |  | 7.47 | 64 | -20 | 0 |
| Left superior temporal gyrus | 9,664 | 7.87 | -60 | -10 | 0 |
| Left superior temporal gyrus |  | 7.83 | -64 | -24 | 2 |

**Table 5.** Peak activations for attentive listening condition greater than noise, young > older adults

| Region | Size (μl) | t score | Coordinates | | |
| --- | --- | --- | --- | --- | --- |
|  |  |  | x | y | z |
| Left superior temporal gyrus | 8,472 | 6.29 | -62 | -16 | 8 |
| Left Heschl's gyrus |  | 4.18 | -40 | -30 | 10 |
| Left Heschl's gyrus |  | 4.08 | -42 | -24 | 12 |
| Right superior temporal gyrus | 3,400 | 4.84 | 52 | -16 | 10 |
| Right superior temporal gyrus |  | 3.37 | 62 | 8 | 2 |

In addition to the overall pattern associated with word perception, we examined psycholinguistic effects of word frequency and phonological neighborhood density using a parametric modulation analysis. There were no significant effects of either word frequency or neighborhood density in the attentive listening condition.

Activity for word perception in the repetition condition (relative to a noise baseline) is shown in **Figure 4**(with maxima in **Tables 6–8**). Again, both young and older adults showed significant activity in bilateral temporal cortex, as well as frontal regions related to articulatory planning, including premotor cortex and supplemental motor area. As with the attentive listening condition, young adults showed significantly more activity in superior temporal regions near auditory cortex. There were no regions where older adults showed more activity than the young adults.

**Figure 4.**
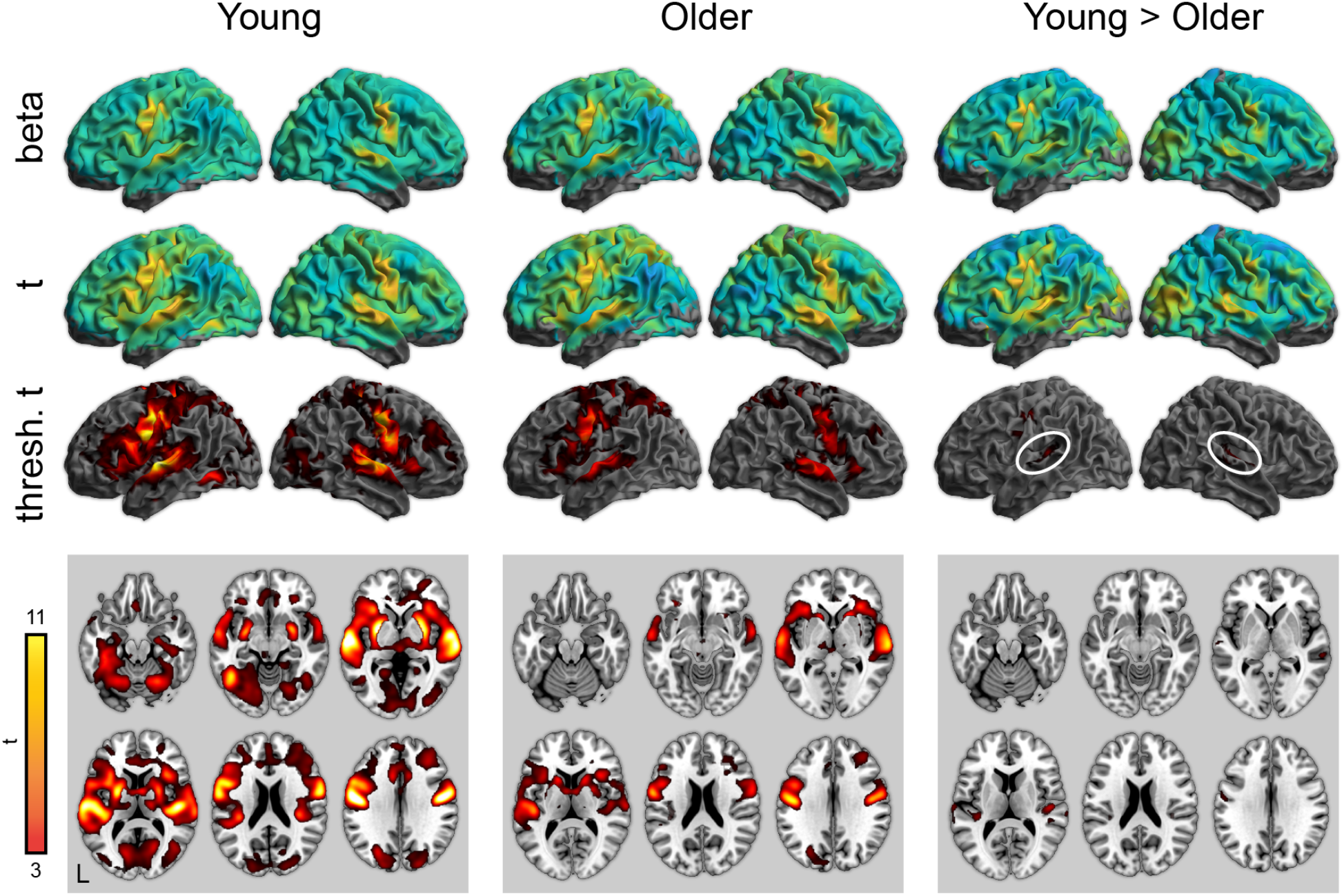
Whole-brain activity for the repetition condition (correct responses only). **Top:** Unthresholded parameter estimates. **Middle:** Unthresholded t maps. **Bottom:** Thresholded t maps (p < .05, cluster corrected). White ovals highlight left and right auditory cortex.

**Table 6.** Peak activations for repetition condition greater than noise, young adults

| Region | Size (μl) | t score | Coordinates | | |
| --- | --- | --- | --- | --- | --- |
|  |  |  | x | y | z |
| Left superior temporal gyrus | 581,432 | 14 | -60 | -14 | 4 |
| Left postcentral gyrus |  | 13.9 | -42 | -16 | 38 |
| Left postcentral gyrus |  | 13.8 | -48 | -14 | 40 |
| Right postcentral gyrus |  | 13.7 | 44 | -12 | 36 |
| Left postcentral gyrus |  | 13.4 | -52 | -8 | 30 |
| Left putamen |  | 12.9 | -24 | 0 | 4 |
| Right superior temporal gyrus |  | 12.5 | 56 | -10 | 4 |
| Supplemental motor area |  | 12.4 | 0 | 0 | 58 |
| Right superior temporal gyrus |  | 12.3 | 52 | -16 | 6 |
| Right superior temporal gyrus |  | 11.4 | 66 | -20 | 2 |
| Right putamen |  | 10.6 | 28 | 0 | -4 |
| Right precentral gyrus |  | 10.6 | 20 | -28 | 60 |
| Left paracentral lobule |  | 10.5 | -18 | -30 | 60 |
| Left Heschl's gyrus |  | 9.89 | -36 | -30 | 14 |
| Left inferior frontal gyrus |  | 9.58 | -52 | 10 | 22 |
| Left inferior temporal gyrus |  | 9.13 | -44 | -56 | -10 |
| Left insula |  | 8.84 | -32 | 22 | 4 |
| Left inferior parietal cortex |  | 8.41 | -38 | -36 | 42 |
| Right insula |  | 8.19 | 34 | 20 | 6 |
| Dorsal anterior cingulate |  | 8.11 | -8 | 12 | 38 |

**Table 7.**
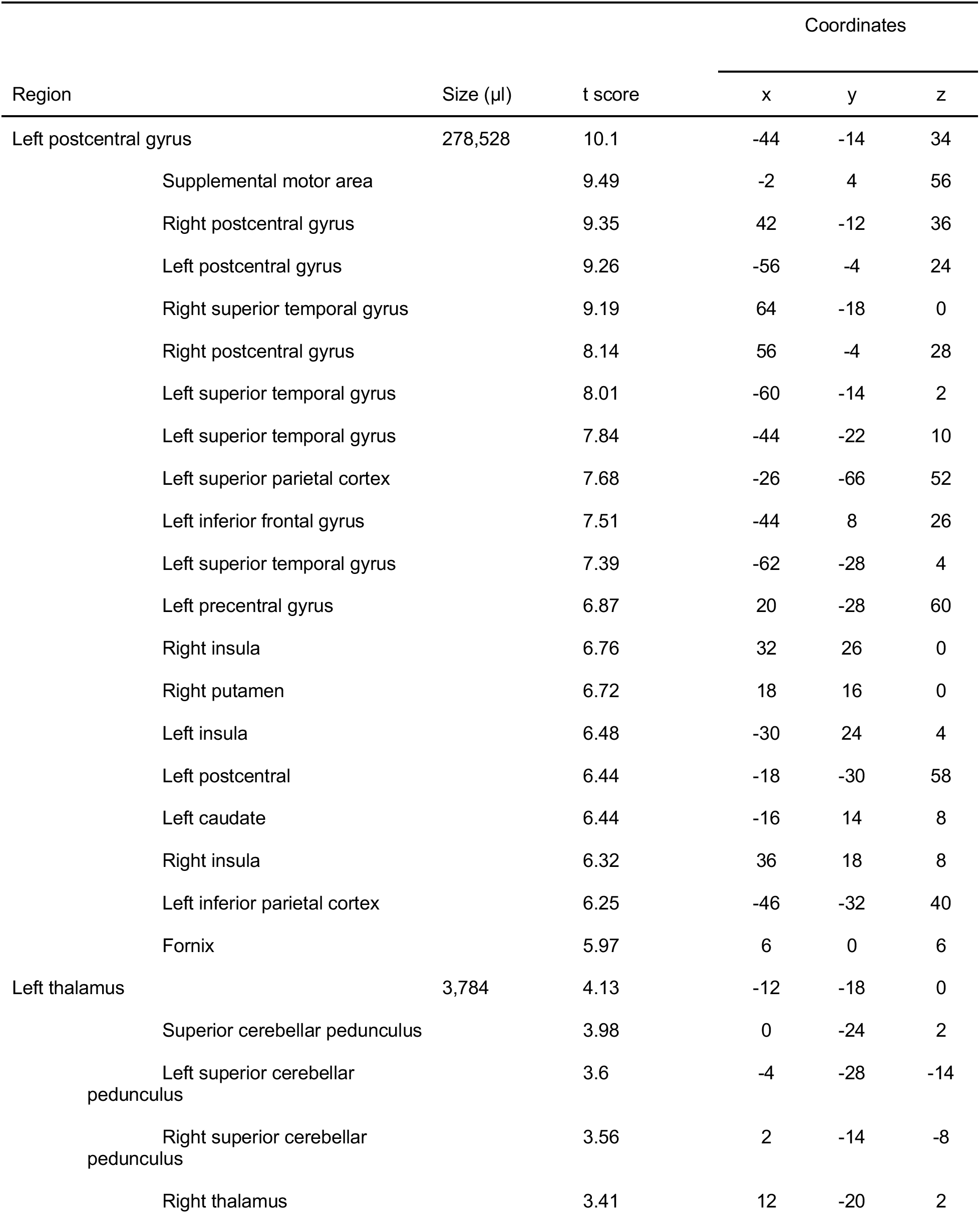
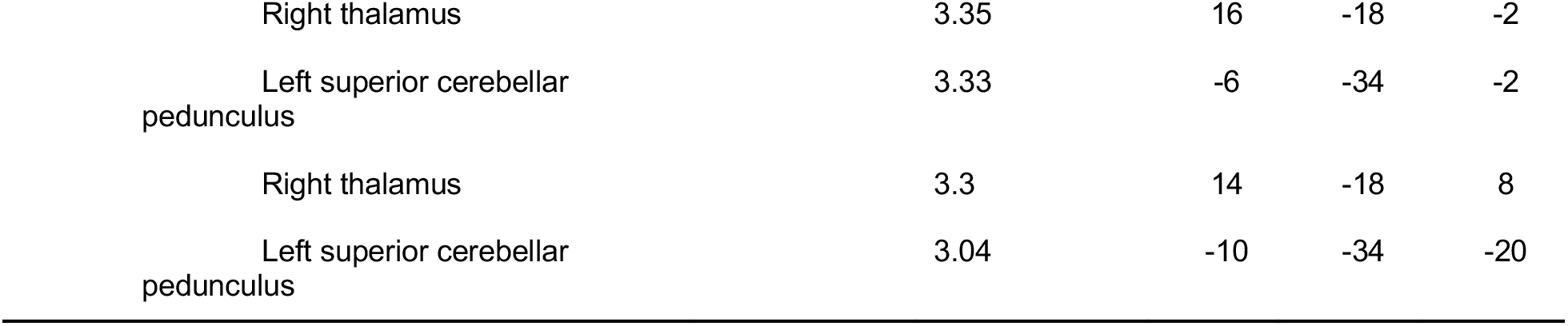
Peak activations for repetition condition greater than noise, older adults

**Table 8.** Peak activations for repetition condition greater than noise, young > older adults

| Region | Size (μl) | t score | Coordinates |  |  |
| --- | --- | --- | --- | --- | --- |
|  |  |  | x | y | z |
| Right Heschl's gyrus | 5,224 | 5.22 | 48 | -20 | 10 |
| Right Heschl's gyrus |  | 4.99 | 40 | -26 | 16 |
| Right superior temporal sulcus |  | 4.18 | 60 | -30 | 0 |
| Right superior temporal gyrus |  | 4.08 | 68 | -26 | 8 |
| Right superior temporal gyrus |  | 3.79 | 56 | -8 | 4 |
| Right superior temporal gyrus |  | 3.62 | 48 | -34 | 10 |
| Left Heschl's gyrus | 4,600 | 4.92 | -36 | -30 | 14 |
| Left superior temporal gyrus |  | 4.17 | -64 | -18 | 8 |
| Left superior temporal gyrus |  | 3.78 | -62 | -32 | 14 |
| Left superior temporal gyrus |  | 3.66 | -60 | -18 | -4 |
| Left Heschl's gyrus |  | 3.47 | -46 | -24 | 6 |
| Left superior temporal gyrus |  | 3.29 | -62 | -38 | 8 |
| Left superior temporal gyrus |  | 3.19 | -52 | -16 | 2 |
| Left postcentral gyrus | 4,248 | 5.26 | -50 | -14 | 44 |
| Left postcentral gyrus |  | 5.21 | -46 | -16 | 42 |
| Left postcentral gyrus |  | 5.12 | -42 | -18 | 40 |
| Left postcentral gyrus |  | 3.88 | -54 | -8 | 28 |

In addition to the overall pattern associated with word perception, we examined psycholinguistic effects of word frequency and phonological neighborhood density using a parametric modulation analysis. There were no significant effects of either word frequency or neighborhood density in the repetition condition.

Finally, we directly compared the attentive listening and repetition conditions, shown in **Figure 5**(with maxima in **Tables 9–10**). Compared to the attentive listening condition, during the repetition condition both young and older listeners showed increased activity in motor and premotor cortex. There were no significant differences between young and older adults.

**Figure 5.**
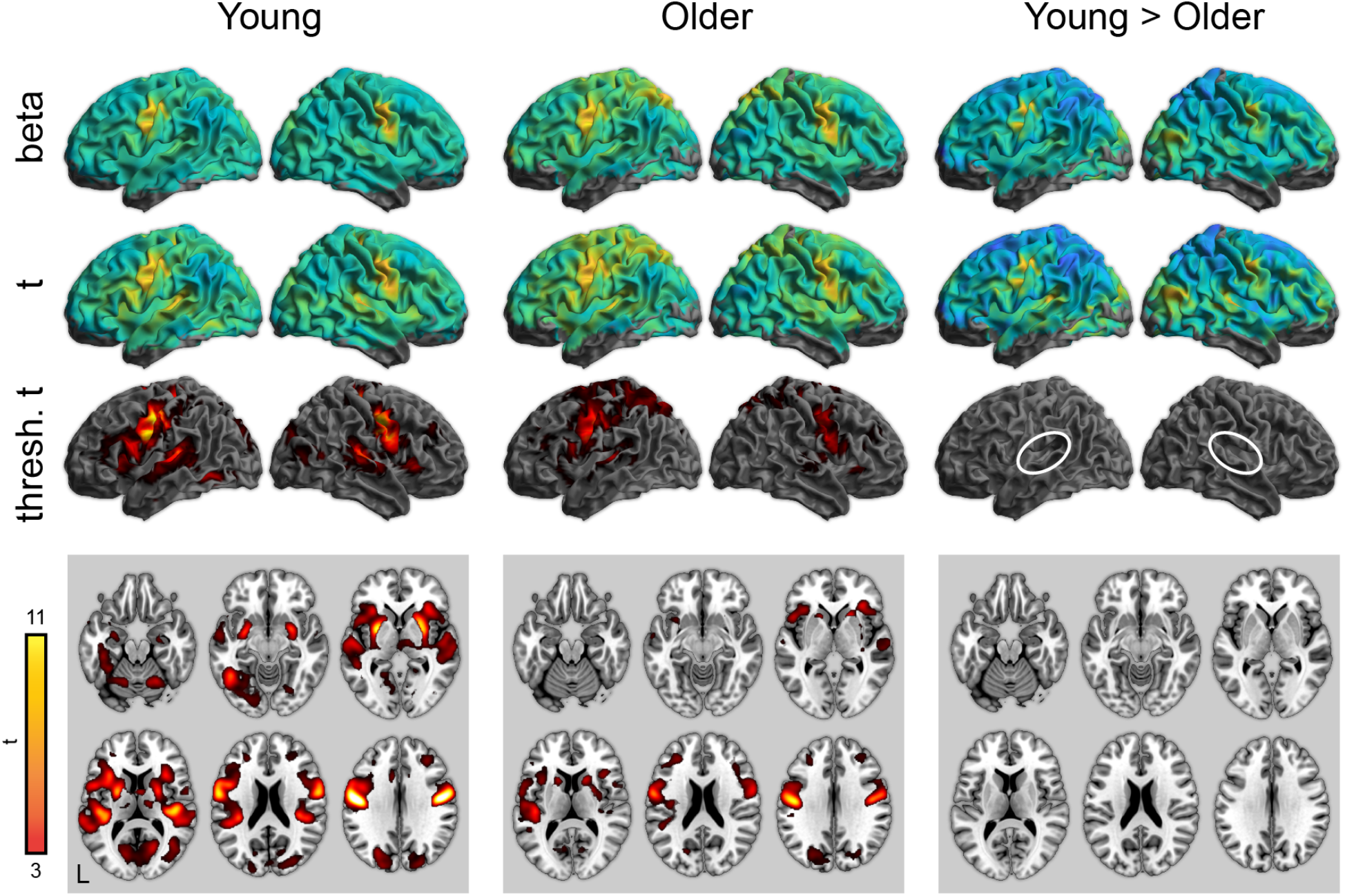
Whole-brain activity for the repetition condition > attentive listening. **Top:** Unthresholded parameter estimates. **Middle:** Unthresholded t maps. **Bottom:** Thresholded t maps (p < .05, cluster corrected). White ovals highlight left and right auditory cortex. There were no significant differences between young and older adults in the repetition > listening contrast.

**Table 9.** Peak activations for word recognition in the repetition condition greater than listening condition, young adults

| Region | Size (μl) | t score | Coordinates | | |
| --- | --- | --- | --- | --- | --- |
|  |  |  | x | y | z |
| Left postcentral gyrus | 311,408 | 13.8 | -44 | -14 | 36 |
| Right postcentral gyrus |  | 13.8 | 44 | -12 | 36 |
| Left putamen |  | 10.4 | -24 | 0 | 4 |
| Supplemental motor area |  | 9.97 | 0 | -2 | 60 |
| Right postcentral gyrus |  | 9.81 | 20 | -28 | 60 |
| Left Heschl's gyrus |  | 9.36 | -36 | -32 | 14 |
| Right superior temporal gyrus |  | 9.11 | 46 | -20 | 8 |
| Left postcentral gyrus |  | 9.06 | -18 | -30 | 60 |
| Right Heschl's gyrus |  | 8.96 | 38 | -26 | 14 |
| Right putamen |  | 8.91 | 28 | 0 | -4 |
| Left insula |  | 7.16 | -32 | 20 | 8 |
| Left inferior frontal gyrus |  | 7.13 | -46 | 8 | 20 |
| Right insula |  | 7.11 | 34 | 22 | 4 |
| Left superior temporal sulcus |  | 6.23 | -52 | -42 | 6 |
| Anterior cingulate |  | 6 | -8 | 12 | 38 |
| Right superior temporal gyrus |  | 5.61 | 64 | -24 | 4 |
| Left inferior parietal cortex |  | 5.6 | -38 | -36 | 42 |
| Right calcarine sulcus |  | 5.47 | 18 | -66 | 8 |
| Right cerebellum |  | 5.31 | 24 | -60 | -22 |

**Table 10.** Peak activations for word recognition in the repetition condition greater than listening condition, older adults

| Region | Size (μl) | t score | Coordinates | | |
| --- | --- | --- | --- | --- | --- |
|  |  |  | x | y | z |
| Left postcentral gyrus | 238,712 | 10.2 | -44 | -14 | 34 |
| Right postcentral gyrus |  | 9.04 | 42 | -12 | 36 |
| Supplemental motor area |  | 7.56 | -2 | 2 | 56 |
| Right postcentral gyrus |  | 7.43 | 54 | -4 | 26 |
| Left Heschl's gyrus |  | 6.45 | -42 | -24 | 10 |
| Left superior parietal cortex |  | 6.43 | -28 | -64 | 54 |
| Left postcentral gyrus |  | 6.03 | -18 | -30 | 60 |
| Right precentral gyrus |  | 6.02 | 20 | -28 | 60 |
| Left precentral gyrus |  | 6 | -46 | 2 | 38 |
| Right putamen |  | 5.88 | 20 | 16 | 0 |
| Right caudate |  | 5.84 | 18 | 14 | 6 |
| Right insula |  | 5.73 | 34 | 28 | 0 |
| Left inferior parietal cortex |  | 5.69 | -44 | -32 | 40 |
| Anterior cingulate |  | 5.49 | -6 | 14 | 42 |
| Left postcentral gyrus |  | 5.44 | -26 | -40 | 62 |
| Left insula |  | 5.38 | -46 | 14 | -2 |
| Right superior parietal cortex |  | 5.29 | 18 | -68 | 56 |
| Left insula |  | 5.28 | -32 | 24 | 4 |
| Left caudate |  | 5.16 | -14 | 14 | 8 |
| Left precentral gyrus |  | 5.13 | -32 | -2 | 64 |
| Right superior temporal gyrus | 4,632 | 4.12 | 58 | -22 | 0 |
| Right Heschl's gyrus |  | 4.11 | 34 | -24 | 4 |
| Right superior temporal gyrus |  | 4.1 | 64 | -16 | 0 |
| Right superior temporal gyrus |  | 4.09 | 52 | -16 | 4 |
| Right posterior insula |  | 3.14 | 32 | -24 | 16 |

## Discussion

We used fMRI to examine neural activity during spoken word recognition in quiet for young and older adult listeners. In both region-of-interest and whole-brain analyses, we found converging evidence for reduced activity in auditory cortex for the older adults. The age differences in auditory cortex activation were present in both an attentive listening task and a word repetition task: Although the repetition task resulted in more widespread activation overall, patterns of age-related differences in auditory cortex were comparable.

There are a number of possible explanations for older adults’ reduced activity during spoken word recognition. One possibility is that age differences in intelligibility might play a role. Intelligible speech is associated with increased activity in a broad network of frontal and temporal regions (Davis and Johnsrude, 2003; Kuchinsky et al., 2012), and in prior studies of older adults, intelligibility has correlated with auditory cortex activity (Harris et al., 2009). We restricted our analyses to correct responses in the repetition condition, and found no statistical support for a relationship between intelligibility and auditory cortex activation (although numerically, participants with better accuracy showed more activity than participants with worse accuracy).

The fact that young and older adults showed comparable activity in auditory cortex during noise trials, with age differences emerging for word recognition trials, is significant. Group differences in activation could be driven not only by neural processing, but also factors such as neurovascular coupling, goodness-of-fit of a canonical hemodynamic response, or movement within the scanner—in other words, artifacts that might differentially impact model parameter estimates in young and older adults but are not of theoretical interest in this context. Although impossible to completely rule out, the selective age differences for speech (but not noise) are consistent with a condition-specific—and thus we argue, neural—interpretation.

Recent evidence suggests age-related changes in temporal sensitivity in auditory regions can be detected with fMRI (Erb et al., 2020). Although our current stimuli do not allow us to explore specific acoustic features, one possibility is that the age-related differences in auditory activity we observed reflect well-known changes in auditory cortical processing that occur in normal aging (Peelle and Wingfield, 2016). Given the increased acoustic complexity of the words relative to noise, acoustic processing differences might drive overall response differences. Such changes may also reflect decreased stimulation as a result of hearing loss; we had insufficient data to rule out this possibility. It is important to note that we cannot completely rule out audibility effects. Even though we limited our responses to correct identification trials, specific acoustic features may still have been less audible for the older adults. It remains an open question whether varying the presentation level of the stimuli would change the age effects we observed.

Age differences in auditory processing are not the only explanation for our results. Auditory cortex is positioned in a hierarchy of speech processing regions that include both ascending and descending projections (Davis and Johnsrude, 2007; Peelle et al., 2010a). Auditory cortex is sensitive not only to changes in acoustic information, but also reflects top-down effects of expectation and prediction (Sohoglu et al., 2012; Wild et al., 2012; Signoret et al., 2018). Thus, the observed age differences in auditory cortex may reflect differential top-down modulation of auditory activity in young and older adult listeners.

Indeed, prior to conducting this study, we expected to observe increased activity (e.g., in prefrontal cortex) for older adults relative to young adults, reflecting top-down compensation for reduced auditory sensitivity. Such activity would be consistent with increased cognitive demand during speech perception in listeners with hearing loss or other acoustic challenge (Pichora-Fuller et al., 2016; Peelle, 2018). Although we were somewhat surprised not to see this, in retrospect, perhaps it would be expected. The stimuli in the current study were presented in quiet, and thus may not have challenged perception sufficiently to robustly engage frontal brain networks. We conclude that during perception of acoustically clear words, older adults do not seem to require additional resources from frontal cortex; whether this changes with increasing speech demands (either acoustic or linguistic) remains an open question.

We did not observe significant effects of either word frequency or phonological neighborhood density on activity during spoken word recognition. These results stand in contrast to prior studies showing frequency effects in visual word perception in fMRI (Kronbichler et al., 2004; Hauk et al., 2008), and word frequency effects in electrophysiological responses (Embick et al., 2001). Prior fMRI studies of lexical competition (including phonological neighborhood density) have been mixed, with some studies finding effects (Zhuang et al., 2011) and others not (Binder et al., 2003). It could be that a wider range of frequency or density or a greater number of stimuli would be needed to identify such effects.

Finally, we found largely comparable age differences in the attentive listening and repetition conditions in auditory cortex. The similarity of the results suggests that using a repetition task may be a reasonable choice in studies of spoken word recognition: although repetition tasks necessarily engage regions related to articulation and hearing one’s own voice, in our data these were not differentially affected by age. An advantage of using a repetition task, of course, is that trial-by-trial accuracy measures can be obtained, which are frequently useful. It is worth noting that our finding of comparable activity in young and older adults for attentive listening and repetition tasks may not generalize to other stimuli or tasks (Davis et al., 2014; Campbell et al., 2016).

A significant limitation of our current study is that we only collected hearing sensitivity data on a minority of our participants. Thus, although we saw a trend towards poorer hearing being associated with reduced auditory cortex activation, it is challenging to draw any firm conclusions regarding the relationship between hearing sensitivity and brain activity. Prior studies using sentence-level materials have found relationships between hearing sensitivity and brain activity in both young (Lee et al., 2018) and older (Peelle et al., 2011) adults. Future investigations with a larger sample of participants with hearing data will be needed to further explore the effects of hearing in spoken word recognition.

From a broader perspective, the link between spoken word recognition and everyday communication is not always straightforward. Much of our everyday communication occurs in the context of semantically meaningful, coherent sentences, frequently with the added availability of visual speech and gesture cues. Given potential age differences in reliance on many of these cues — including older adults’ seemingly greater reliance on semantic context (Rogers 2016; Rogers et al. 2012; Wingfield and Lindfield 1995)— it seems likely that our findings using isolated spoken words cannot be extrapolated to richer naturalistic settings.

In summary, we observed largely overlapping brain regions supporting spoken word recognition in young and older adults in the absence of background noise. Older adults showed less activity than young adults in auditory cortex when listening to words, but not noise. These patterns of age difference were present regardless of the task (attentive listening vs. repetition).

## Acknowledgements

The multiband echo planar imaging sequence was provided by the University of Minnesota Center for Magnetic Resonance Research. We are grateful to Linda Hood for assistance with data collection, and to Henry Greenstein, Ben Muller, Olivia Murray, Connor Perkins, and Tracy Zhang for help with data scoring.

## Footnotes

1 The room was not sound isolated and low frequency noise from building HVAC was typically present.

2 The flip angle was suboptimal due to an error setting up the sequences; although discovered partway through the study, we left it unchanged to maintain consistent data quality. With a TR of ~ 3 seconds we would expect a better signal-to-noise ratio with a flip angle of 90°.

